# Antigenic imprinting dominates humoral responses to new variants of SARS-CoV-2 in a hamster model of COVID-19

**DOI:** 10.1101/2024.10.30.621174

**Authors:** Joran Degryse, Elke Maas, Ria Lassaunière, Katrien Geerts, Yana Kumpanenko, Birgit Weynand, Piet Maes, Johan Neyts, Hendrik Jan Thibaut, Yeranddy A. Alpizar, Kai Dallmeier

## Abstract

The emergence of SARS-CoV-2 variants escaping immunity challenges the efficacy of current vaccines. Here, we investigated humoral recall responses and vaccine-mediated protection in Syrian hamsters immunized with the third-generation Comirnaty® Omicron XBB.1.5-adapted COVID-19 mRNA vaccine, followed by infection with either antigenically closely (EG.5.1) or distantly related (JN.1) Omicron subvariants. Vaccination with YF17D vector encoding a modified Gamma spike (YF-S0*) served as a control for pre-Omicron SARS-CoV-2 immunity. Our results show that both Comirnaty® XBB.1.5 and YF-S0* induce robust, however, poorly cross-reactive, neutralizing antibody (nAb) responses. In either case, total antibody and nAb levels increased following infection. Intriguingly, the specificity of these boosted nAbs did not match the respective challenge virus but was skewed towards the primary antigen used for immunization, suggesting a marked impact of antigenic imprinting; confirmed by antigenic cartography. Furthermore, limited cross-reactivity and rapid decline of nAbs induced by Comirnaty® XBB.1.5 with EG.5.1 and, more concerning, JN.1. raises doubts about sustained vaccine efficacy against recent circulating Omicron subvariants. Future vaccine design may have to address two major issues: (i) to overcome original antigenic sin that limits the breadth of a protective response towards emerging variants, and (ii) to achieve sustained immunity that lasts for at least one season.

## 1. Introduction

The high mutation rate of SARS-CoV-2 raises continuous concerns regarding the sustained efficacy of existing vaccines. Since the onset of the COVID-19 pandemic, which has led to over 776 million cases and more than 7 million deaths globally [1], the virus has rapidly evolved, producing antigenic variants that challenge current vaccination strategies. With the emergence of new variants, there is hence an urgent need to reassess vaccine performance.

The COVID-19 mRNA vaccine Comirnaty® (Pfizer/BioNTech), originally developed to target the ancestral SARS-CoV-2 strain, has undergone several modifications to follow emerging variants, to avoid breakthrough infections in the vaccinated population due to antigenic drift. The most recent, third-generation of Comirnaty® (updated in August 2023), targeting the Omicron subvariant XBB.1.5, was rolled out during infection waves dominated by XBB subvariants (XBB.1.5, EG.5.1), and spanning to the current surge (starting end of 2023) of BA.2.86 subvariants (JN.1, JN.1.16, KP.3) (Figure 1a). BA.2.86 originates from another sublineage of Omicron than XBB and harbours 36 additional mutations in its spike protein compared to XBB.1.5 [2], representing a unique antigenic composition that arose within the BA.2 lineage. Compared to founding BA.2.86, JN.1 acquired one additional mutation on the receptor binding domain (L455S), reducing the affinity for human ACE2 binding. However, this mutation has been shown to greatly enhance resistance to pre-existing humoral immunity, particularly towards neutralizing antibodies (nAbs) established by monovalent XBB.1.5 vaccines or prior infections with XBB.1.5 or EG.5.1 [3, 4].

**Figure 1.**
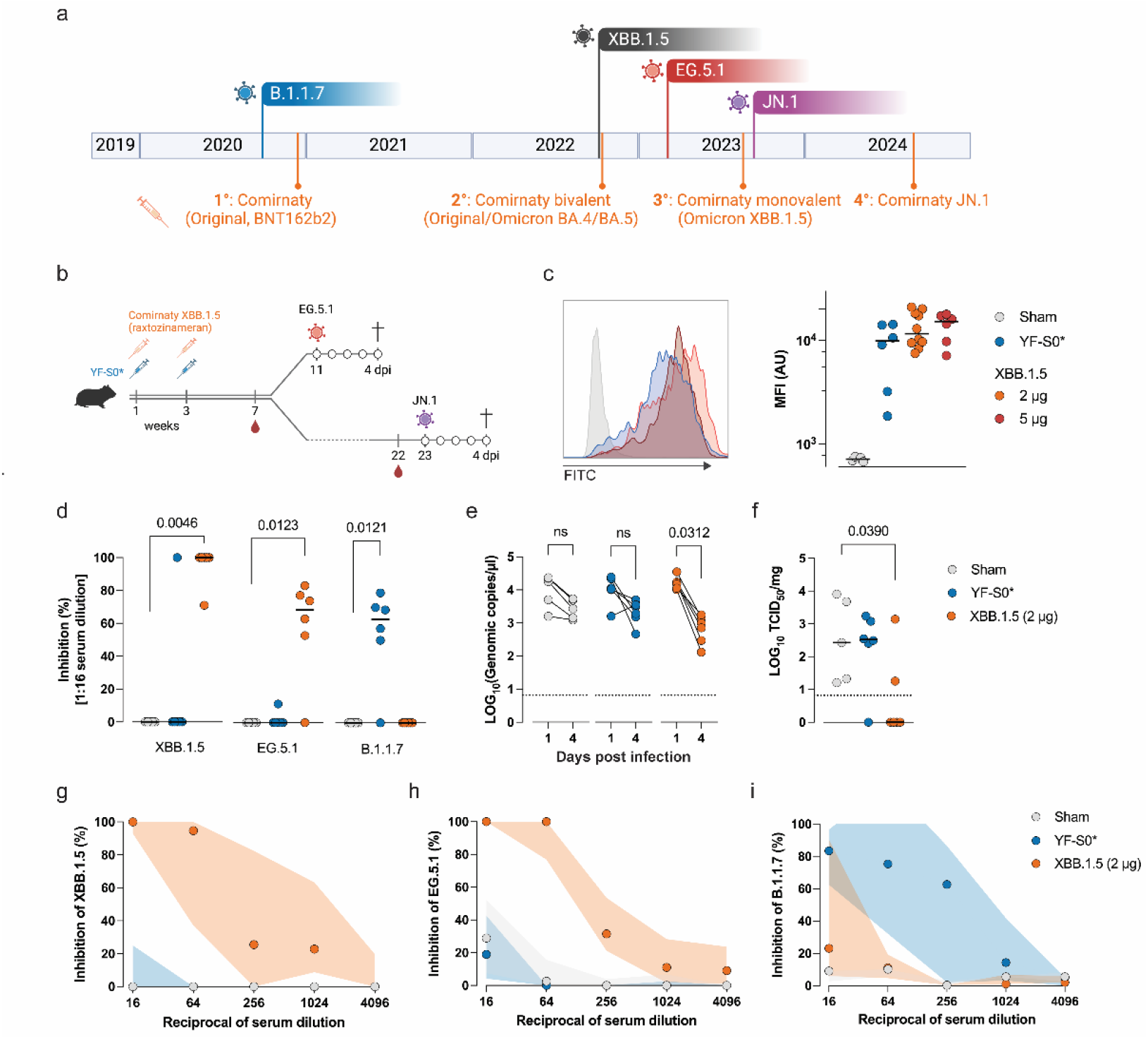
Vaccine-induced neutralizing antibodies against EG.5.1 are boosted after autologous infection. **(a)** Emergence of selected historical (B.1.1.7) and more recent Omicron SARS-CoV-2 variants along with a timeline for the introduction of original (1° generation) and updated (2° to 4°) mRNA vaccines. **(b)** Schematic representation of the experimental design. **(c)** Representative histograms of flow cytometry showing the recognition of spike protein by sera from animals vaccinated with Comirnaty® XBB.1.5 (range and red) and YF-S0* (blue). Sham-vaccinated animals (grey) were used as negative controls. Median fluorescence intensity (MFI) of individual animals in represented on the right panel. Horizontal bars represent median values. **(d)** Inhibition of cytopathic effect by sera in cells infected with XBB.1.5, E.G.5.1 and B.1.1.7. Data points represent individual animals. Horizontal bars represent median values. Dunn’s multiple comparison test. **(e)** Viral loads in throat swabs quantified by qRT-PCR. Paired comparisons using Wilcoxon test. **(f)** Virus titration from lungs samples. Horizontal bars represent median values. Dunn’s multiple comparison test. **(g-i)** Neutralization curves (expressed as relative inhibition) against XBB.1.5 **(g)**, EG.5.1 **(h)** and B.1.1.7 **(i)** from sera of animals challenged with EG.5.1. Data points represent median values (n = 5, 6, 6 for Sham, YF-S0* and XBB.1.5 groups, respectively); shaded areas indicate interquartile ranges (IQR).

In individuals with prior exposure to ancestral SARS-CoV-2, updated SARS-CoV-2 mRNA vaccines effectively recall B cells to produce neutralizing antibodies against conserved epitopes with the ancestral spike protein [5]. Though such an imprinted immune recall may contribute to vaccine protection conferred by variant booster vaccines [6-13], it may limit the de novo generation of variant-specific antibodies that target epitopes associated with immune escape, particularly in antigenically distant variants. Such failure to redirect immunity by another antigen exposure has been coined original antigenic sin, acknowledging a skewed memory response that leads to greater induction of antibodies specific to the first-encountered variant of an immunogen instead of its subsequent variants [14].

Here, we investigate the magnitude and quality of humoral recall responses as well as the degree of vaccine-mediated protection in Syrian hamsters vaccinated with Comirnaty® Omicron XBB.1.5 COVID-19 mRNA vaccine when exposed either to an antigenically close XBB subvariant (EG.5.1) or distant BA.2.86 subvariant (JN.1). We demonstrate that nAbs against the autologous antigen dominates the humoral response upon infection. Unlike total binding antibodies against ancestral spike protein and XBB.1.5-specific nAbs, EG.5.1- and JN.1-specific nAbs wane rapidly and cannot be recovered after infection with heterologous (antigenically mismatched) JN.1. These results underscore the need for continuous vaccine updates to maintain effective protection against evolving SARS-CoV-2 variants and highlight the challenges of addressing antigenic evolution in future vaccination strategies.

## 2. Materials and Methods

### 2.1. Viruses and Vaccines

The following SARS-CoV-2 strains were used in this study: VOCs Alpha (lineage B.1.1.7; hCoV104 19/Belgium/rega-12211513/2020; EPI_ISL_791333), Omicron subvariant XBB.1.5 (EPI_ISL_17273054) Omicron subvariant EG.5.1 (SARS-CoV-2/hu/DK/SSI-H121; OY747654) and Omicron subvariant JN.1 (SARS-CoV-2/hu/DK/SSI-H137, passage 2 stock, isolated from human via nose and throat swab on 26 Dec 2023 in Zealand, Denmark). All work related to these viruses was conducted in the high-containment BSL3 facilities of the KU Leuven Rega Institute (3CAPS) under licenses AMV 30112018 SBB 219 2018 0892 and AMV 23102017 SBB 219 2017 0589 according to institutional guidelines.

Comirnaty® Omicron XBB.1.5 COVID-19 mRNA vaccine vials (Lot number: HD9868, 0.1 µg Raxtozinameran/µl Pfizer/BioNTech, Mainz, Germany), were recovered from vaccination centers and transported on ice. Vaccinations were performed within 12 h after thawing. The YF-S0*-vaccine was produced in-house as previously described [15, 16].

### 2.2. Cells

Vero E6 cells were maintained in minimum essential medium (MEM; Gibco, Thermo Fisher Scientific, Waltham, Massachusetts, USA) supplemented with 10% fetal calf serum (FCS, HyCloneTM, Logan, Utah, USA), 1% Pen Strep Glutamine (Gibco), 1% non-essential amino acids (NEAA, Gibco). A549-hACE2-TMPRSS2 (Invivogen) were kept in 10% FCS DMEM supplemented with puromycin (0.5 µg/ml) and hygromycin (100 µg/ml).

HEK293 stably expressing the SARS-CoV-2 spike-protein (Wuhan, D614G), hereafter referred as Mono8 cells, were provided by TPVC (Translational Platform Virology and Chemotherapy) and kept in DMEM supplemented with 10% FCS (HyCloneTM), 1% Pen Strep (Gibco) and 10 µg/ml Blasticidin (InvivoGen, San Diego, California, USA).

### 2.3. Animals

Male Syrian golden hamsters (Mesocricetus auratus, RjHan: AURA strain; 7–8-week-old), were bred in-house. Housing conditions and experimental procedures were approved by the ethics committee of animal experimentation of KU Leuven (license number: P055-2021), following institutional guidelines approved by the Federation of European Laboratory Animal Science Associations (FELASA).

### 2.4. Syrian golden hamster SARS-CoV-2 infection model

M. auratus as an infection model for SARS-CoV-2 has been previously described [15]. All animals were between 7 and 25 weeks old at beginning of the experiment. Hamsters were vaccinated intramuscularly (i.m.) with 2 µg (n = 10) or 5 µg (n = 6) Raxtozinameran (formulated in 20 µl and 50 µl of Comirnaty®, respectively) on day 0 (d0) and received a booster vaccination of the same dosing on day 20. Control cohorts received 2 doses of: 104 PFU YF-S0*-vaccine [16] (n = 6) or MEM (Gibco) supplemented with 2% FCS (HyCloneTM) (Sham) (n = 5) intraperitoneally (i.p.), on the same days. Blood was drawn via the jugular vein on day 48 and 157 to check for antibodies. Six Comirnaty®-vaccinated (2 µg Raxtozinameran), six YF-S0*-vaccinated animals and five sham-vaccinated animals were intranasally infected with 100 µl, 104 TCID50 EG.5.1, on day 77 (week 11). Four Comirnaty®-vaccinated (2 µg Raxtozinameran) and 5 non-treated age-matched wild type hamsters (25 weeks old), were intranasally infected with 100 µl, 104 TCID50 JN.1, on day 164 (week 22). After infection, throat swabs (CLASSIQSwabs, Copan Italia s.p.a., Brescia, Italy) were collected and animals were weighed daily. Four days post infection (dpi) all hamsters were euthanized for the collection of lung tissue and serum.

### 2.5. SARS-CoV-2 antibody detection with Mono8 cells

Total anti S-protein IgG antibodies induced by vaccination were evaluated in serum samples by flow cytometry using HEK cells stably transfected with spike protein from Wuhan-Hu-1 strain (Mono8 cells). Mono8 cells (105 cells/well) were plated in a U-bottom 96-well plate (Corning Inc., Corning, New York, USA) and blocked from 20 min in ice-cold FACS buffer (2% fetal calf serum in Dulbecco’s Phosphate Buffered Saline (Gibco) supplemented with 2.5 mM EDTA (Invitrogen, Thermo Fisher Scientific, Waltham, Massachusetts, USA)). The cells were pelleted by centrifugation at 500 x g for 2 min and resuspended in prediluted serum samples (1:20 in FACS buffer). Cells were incubated on ice, for 20 min. After two washing steps, the cells were incubated for 20 min on ice with FITC-conjugated rabbit anti-hamster IgG antibody (1:500 in FACS-buffer; 307-095-003, Jackson ImmunoResearch Labs, West Grove, Pennsylvania, USA). Subsequently, the cells were washed and incubated (15 min, RT, protected from light) in a 4% paraformaldehyde solution (PFA; Thermo Fisher Scientific) for 10 min at room temperature, protected from light. After fixation the cells were washed and resuspended in FACS-buffer. Acquisition was performed in a BD LSRFortessa™ X-20 Cell Analyzer (BD Biosciences, Franklin Lakes, New Jersey, USA). Data analysis and representation were performed in FlowJo™ v10.8 Software (BD Life Sciences).

### 2.6. Serum Neutralization Test

To measure SARS-CoV-2 neutralizing antibodies (nAbs), serial dilutions of heat-inactivated serum samples (25 µl) were mixed with an equal volume of 100 TCID50 of SARS-CoV-2 B.1.1.7 and incubated for 1 hour at 37 °C. Following incubation, the serum-virus mixtures were added to 96-well plates containing Vero E6 (B.1.1.7, XBB.1.5, EG.5.1) or A549-hACE2-TMPRSS2 (JN.1) cells that had been seeded the previous day at 104 cells per well. Cell viability was assessed after 96 hours using an MTS reduction assay. To this end, the medium was aspirated from the wells, and 75 µl of an MTS/phenazine methyl sulfate (PMS) solution (2 mg/ml MTS (Promega) and 46 μg/ml PMS (Sigma–Aldrich) in PBS at pH 6–6.5) was added. Following a 2-hour incubation at 37 °C, absorbance was measured at 498 nm with a Spark® multimode microplate reader (Tecan). Serum neutralization titres (SNT50) were defined as the last dilution with detectable signal.

### 2.7. RNA isolation

Hamster lung tissues were collected and homogenized using bead disruption (Precellys kit, Bertin Corp., Rockville, MD, USA, catalog # 432-0141) in TRK lysis buffer (E.Z.N.A.® Total RNA Kit, Omega Bio-tek Inc, Norcross, GA, USA, catalog # R6834-02) and centrifuged (10,000 rpm, 5 min) to pellet the cell debris. The NucleoSpin kit (Macherey-Nagel) was used for throat swabs. RNA was extracted according to the manufacturer’s instructions and eluted in 50 μl. From this, 4 μl were use as template for RT-qPCR reactions. RT-qPCR was performed using a LightCycler® 96 and iTaq Universal Probes One-Step RT-qPCR kit (BioRad, Temse, Belgium), with purified RNA and primers and probes specific to SARS-CoV-2 as reported by Boudewijns et al. [17] SARS-CoV-2 cDNA control (Integrated DNA Technologies, Leuven, Belgium) was used to calculate genome copies per mg tissue or ml of oral swab. Cytokine expression levels were normalized to the mean expression β-actin transcripts. The lower limit of detection (LLOD) for SARS-CoV-2 was estimated at 7150 copies per ml, based on analysis of repeated standard curves. Samples with values below the LLOD were considered negative for expression and assigned the same value.

### 2.8. Endpoint titrations

The lung tissues were homogenized using bead disruption (Precellys kit, Bertin Corp., Rockville, MD, USA) in 350 µl of minimal essential medium. After centrifugation (10,000 rpm, 5 min, 4 °C) to pellet the cell debris, endpoint titrations in triplicates were carried out on confluent Vero E6 cells seeded on 96-well plates. Viral titers were determined using the Reed and Muench method with the Lindenbach calculator and were expressed as 50% tissue culture infectious dose (TCID50) per mg tissue.

### 2.9. Antigenic cartography

Antigenic cartography was performed in RStudio (Version 2024.04.2+764). Maps were computed using the Racmacs package (https://acorg.github.io/Racmacs/, version 1.1.35) [16, 18], using the reciprocal of the last neutralizing titre and 500 optimizations. The minimum column basis was set to “none”. Neutralizing titres below the limit of detection were set to 1 for all calculations.

## 3. Results

### 3.1. Evaluation of vaccine efficacy of Comirnaty® XBB.1.5 against EG.5.1

To assess the contribution of antigenic imprinting to the recall response to EG.5.1, immunocompetent Syrian hamsters were vaccinated with either of two vaccines encoding antigenically distant spike proteins, namely Raxtozinameran (Comirnaty® XBB.1.5) or YF-S0* [16]. Comirnaty® XBB.1.5 (2 µg or 5 µg) was administered intramuscularly twice, three weeks apart (Figure 1b), emulating the two-dose regime followed in humans for primary immunisation. A separate group of hamsters received two intraperitoneal administrations of YF-S0* (104 PFU), three weeks apart. YF-S0* is a second-generation YF17D-vectored vaccine candidate updated (in 2021) to encode a prefusion stabilized spike protein of more ancestral VOC Gamma (P.1) [15, 16]. Seven weeks after vaccination, we confirmed that 100% of Comirnaty® XBB.1.5 vaccinated hamsters had seroconverted, with high IgG antibody titres cross-reactive to ancestral Wuhan (B1.1 D614G lineage). At this time point, we did not observe significant differences in total IgG responses induced by 2 µg (MFI 4.1 log10, 95% CI 3.9-4.3) and 5 µg (MFI 4.2 log10, 95% CI 3.8-4.3) doses of XBB.1.5. Live-attenuated YF-S0* elicited equally high anti-S IgG levels (MFI 3.9 log10, 95% CI 3.2-4.2). As expected, anti-S antibodies were not detected in sham-vaccinated animals (Figure 1c).

Next, we evaluated the levels of nAbs against the autologous XBB.1.5 and the closely-related XBB subvariant EG.5.1. VOC Alpha (B.1.1.7) was used as a control to assess crossreactive nAbs induced by Omicron-based Comirnaty® XBB.1.5 and pre-Omicron YF-S0*. Two 2-µg-doses of Comirnaty® XBB.1.5 induced high titres of nAbs against XBB.1.5 (GMT 256, 95% CI 32.7-2003). Accordingly, at the highest serum concentration tested (1:16), these nAbs were capable of fully inhibiting the cytopathic effect in infected cells (Figure 1d). Likewise, nAbs against EG.5.1 were detected in 5/6 XBB.1.5-vaccinated animals at the highest serum concentration (GMT 20.2, 95% CI 2.9-139.4) with a median inhibition of CPE of 64%. However, their neutralizing activity dropped to undetectable levels in most cases already in the subsequent serum dilution (data not shown). By contrast, only 1/6 YF-S0*-vaccinated hamsters had detectable XBB.1.5 nAbs and only at the lowest serum dilution tested (GMT 1.6, 95% CI 0.5-5.2) (Figure 1d). In contrast, vaccination with XBB.1.5 did not induce nAbs against B.1.1.7. We have previously shown that YF-S0* induces robust nAb responses against pre-Omicron variants as well as original Omicron variant (B.1.1.529; from 12/2021), yet with lower potency [16]. Accordingly, YF-S0* vaccination elicited nAbs against B.1.1.7 (GMT 32, 95% CI 4.3-237.7), but failed to induce marked nAbs against the more recent Omicron subvariants XBB.1.5 and EG.5.1 (GMT 1.6, 95% CI 0.5-5.2). Thus, in this simplified immunization model where naïve animals were exposed each to a single monovalent vaccine antigen, we observed a robust cross-reactive binding antibody response, but no cross-reactive nAbs. In this setting, we sought to trigger recall responses by intranasal challenge with EG.5.1.

Vaccinated hamsters and sham-injected controls were inoculated with 104 TCID50 of EG.5.1 and sacrificed 4 days post-infection. Body weight measurements and throat swabs were performed daily. In line with previous reports [19], EG.5.1 infection was mild and did not affect total body weight in any of the experimental groups (data not shown). Viral replication in upper airways (longitudinally) and in the lungs (at endpoint) was determined by qPCR and virus titration, respectively. Quantification of viral RNA loads in throat swabs revealed a slight yet a not significant decay of viral transcripts in both sham-injected and YF-S0*-vaccinated animals over the course of infection, possibly associated with natural clearance (Figure 1e). EG.5.1 RNA was also still detected in animals immunized with the more recent XBB.1.5 vaccine 4 days post infection yet at significantly lower levels and prone to faster elimination (10.4-fold reduction, p = 0.0312, paired t test). Importantly, in 4/6 XBB.1.5-vaccinated hamsters infectious virus in the lungs was reduced to undetectable levels (Figure 1f). Of note, the only XBB.1.5-vaccinated animal with high viral titres in the lungs corresponds to that lacking EG.5.1-specific nAbs (Figure 1d). The outdated, VOC-based YF-S0* candidate did not reduce infectious particle counts in the lung, except for one animal with low, but detectable, EG.5.1 nAbs prior to challenge.

Serum neutralization assays using post challenge sera revealed a 10-100-fold increase in XBB.1.5 nAbs after EG.5.1 challenge (Figure 1g, GMT 1625.5, 95% CI 495-5332). A much larger increase was observed in EG.5.1-specific nAbs (103-104-fold, GMT 256, 95% CI 40.6-1612), suggesting that the rise in XBB.1.5 nAbs levels likely corresponded to a recall response of cross-reactive B cells against EG.5.1. In contrast, only a poor neutralization effect against B.1.1.7 was observed, i.e. in 3/6 animals and then at the lowest serum dilution (Figure 1g-i). In YF-S0*-vaccinated animals, XBB.1.5- and EG.5.1-nAbs were barely detectable in 3/6 and 1/6 hamsters at the highest concentration of serum, respectively, suggesting that cross-reactive B cells against pre-Omicron variants (i.e. B.1.1.7) might have been induced, though at very low frequencies. However, nAb levels as tested against B.1.1.7 were significantly boosted (16-fold change, GMT 128, 95% CI 8.4-1946.0) despite the lack of cross-reactive nAbs prior to the challenge. These results indicate that in the absence of pre-exposure to ancestral SARS-CoV-2, new antigenically distant variants fail to effectively trigger cross-reactive nAbs, despite robust levels of cross-reactive binding antibodies. Hence, with respect to nAbs, old and new variants appear to behave like distinct serotypes. In addition, our findings strongly suggest that antigenic imprinting by previous encounter (in this case, by vaccination) dominates the subsequent humoral response to new SARS-CoV-2 variants.

### 3.2. Evaluation of the long-term efficacy of Comirnaty® XBB.1.5 against antigenically distant variants

Next, we assessed the longevity of the humoral response induced by Comirnaty® XBB.1.5 using sera collected 22 weeks after vaccination (Figure 1b). Binding antibody titres decreased significantly (e.g. 4.9 log10 at 1:100 dilution, p < 0.001) compared to 7 weeks after the first vaccination; but remaining at relatively high levels compared to sham-vaccinated controls (Figure 2a). XBB.1.5 nAbs were still detectable after 22 weeks in 3/4 hamsters (GMT 32, 95% CI 0.4-2186). However, EG.5.1 nAbs had dropped to undetectable levels in 3/4 hamsters (GMT 2.0, 95% CI 0.2-18.2, Figure 2b). In addition, we evaluated whether nAbs induced by XBB.1.5 still covered the novel antigenically distant JN.1 variant, which had become the prevalent circulating strain among the American and European populations at the time these hamsters had been vaccinated for 22 weeks (in February 2024) [20, 21] (Figure 1a). Seven weeks after vaccination with Comirnaty® XBB.1.5, 3/4 hamsters had JN.1 nAbs, but at much lower levels than EG.5.1 nAbs (GMT 8.0, 95% CI 0.8-72.6). JN.1 nAbs had waned to undetectable levels in all hamsters by week 22 (Figure 2b).

**Figure 2.**
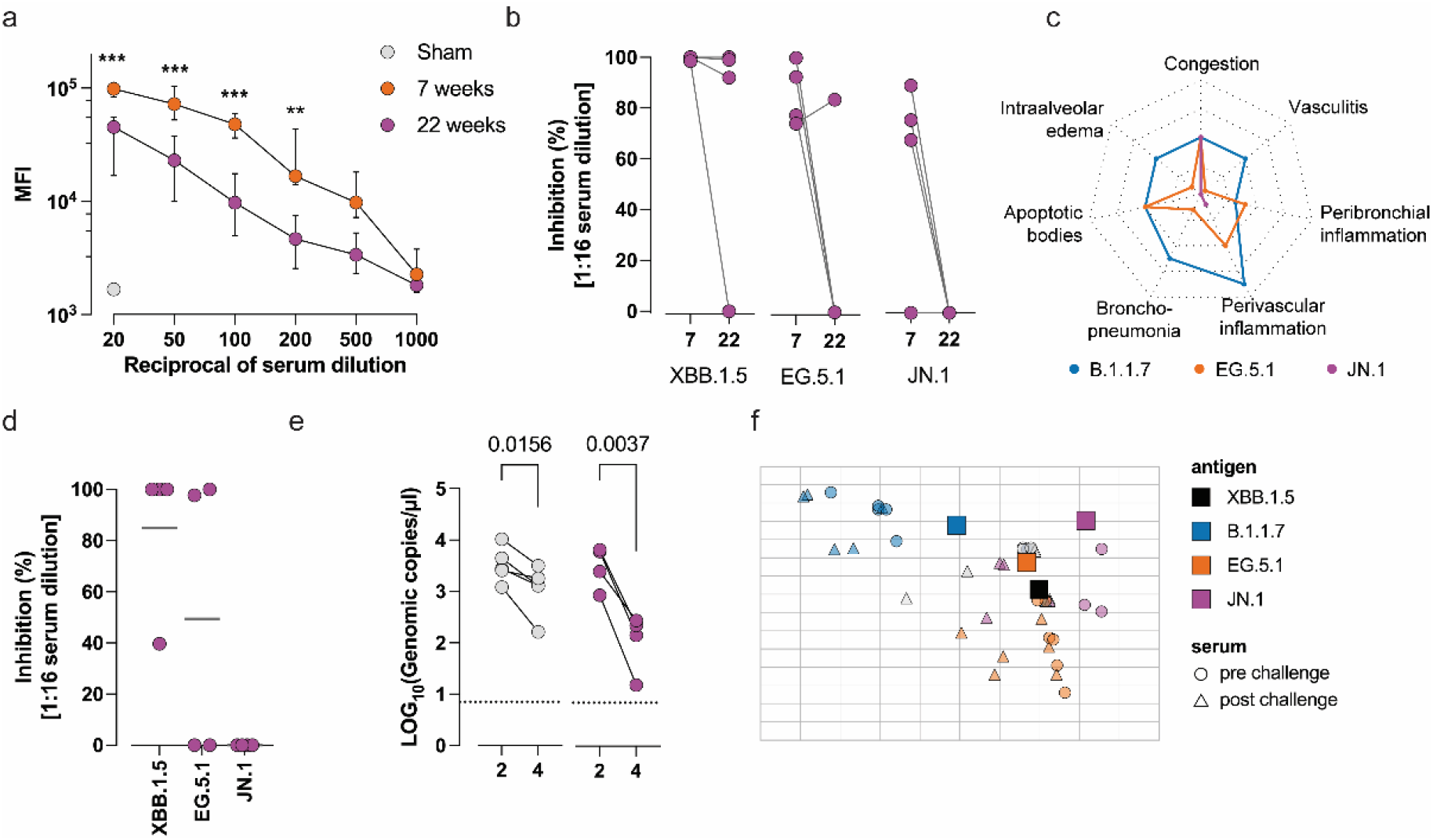
Waning neutralizing antibodies are not rescued after challenge with JN.1. **(a)** Median fluorescence intensity (MFI) indicative for anti-SARS-CoV-2 binding IgG from serum from Comirnaty® XBB.1.5-vaccinated animals at 7 and 22 weeks after vaccination (as in Figure 1c). The highest dilution of serum from sham-vaccinated animals is included as negative control. Data points represent the median ± IQR. ***, p<0.001; p<0.01; Šidak’s multiple comparison test, with a single pooled variance. **(b)** Neutralization of XBB.1.5, EG.5.1 and JN.1 variants by serum from XBB.1.5-vaccinated animals at weeks 7 and 22 after first vaccination. **(c)** Mean lung pathology score at 4 days post infection with B.1.1.7, EG.5.1 and JN.1. Score range 0-2 (normal-severe), each segment indicates 0.5 score units. **(d)** Neutralization of XBB.1.5, EG.5.1 and JN.1 by 4 days post infection sera from animals challenged with JN.1. Horizontal bars represent median values (n = 4) **(e)** Viral loads in buccal swabs quantified by qRT-PCR. Paired comparisons using Wilcoxon test. **(f)** Antigenic cartography. Two-dimensional space representing the cross-reactivity of nAbs from pre-challenge sera (circles) raised by XBB.1.5 (orange, purple) and YF-S0* (blue) vaccine antigen against SARS-CoV-2 variants (squares, indicated as antigens). Sera from Sham-vaccinated animals are represented in grey. Triangles denote serum nAbs after challenge with EG.5.1 (orange, blue) and JN.1 (purple). Each square in the grid represents one antigenic distance unit.

Next, we infected the vaccinated hamsters intranasally with JN.1 (104 TCID50). A cohort of naïve hamsters were infected with B.1.1.7 and used as experimental benchmark for severe lung pathology [22]. Similar to its parental variant BA.2.86 [19], JN.1 infection did not evoke any change in body weight (data not shown) and caused a milder lung pathology compared to EG.5.1 with very low to undetectable peribronchial/perivascular inflammation at 4 dpi (Figure 2c). Recall responses in XBB.1.5-vaccinated hamsters after JN.1 infection were predominantly targeted against the cognate vaccine antigen; namely, a robust increase in XBB.1.5 nAbs in 4/4 hamsters (GMT 32.0, 95% CI 3.5-290.5). NAbs against XBB.1.5 descendent EG.5.1 could be detected at least in 2/4 animals at the highest serum concentration (GMT 4.0, 95% CI 0.3-51.1). By contrast, nAbs against JN.1 remained undetectable (Figure 2d). Taken together, these results suggest that, irrespective of the spike protein expressed by the challenge virus, B cells imprinted by vaccination towards a specific, possibly different, or increasingly mismatched, antigen are preferentially reactivated.

Despite the absence of JN.1-specific nAbs, viral RNA levels in throat swabs decreased significantly by day 4 post-infection, suggesting that other effector mechanisms, such as innate antiviral responses, cellular immunity, or Fc-mediated mechanisms, play a role in viral clearance in the upper airways (Figure 2e). Moreover, as observed before for EG.5.1 (Figure 1e), also in non-vaccinated animals, viral RNA levels in the upper airways showed a significantly decline 4 dpi. Neither infectious particles nor viral transcripts were detected in the lungs at 4 dpi (data not shown), confirming the reduced pathogenicity and propensity of BA.2.86 subvariants to replicate in Syrian hamsters [19].

Antigenic relatedness of the thus studied viral spike variants and resulting immune imprinting was further analyzed using antigenic cartography of nAb responses [16, 18]. In agreement with previous reports, we observed that EG.5.1 clustered closer to its parent XBB.1., while B.1.1.7 (VOC Alpha) and JN.1 (offspring of distinct Omicron sublineage BA.2.86) stand separated from XBB.1.5, each at opposite sides; indicative for a growing antigenic difference along the evolutionary trajectory (Figure 2f, squares). Fully as expected, sera from XBB.1.5 vaccinated animals clustered in close proximity to XBB.1.5, whereas YF-S0* sera grouped next to pre-Omicron VOC Alpha (B.1.1.7). Sera coordinates did not change significantly neither for the XBB.1.5-vaccinated animals (orange triangles) nor for the YF-S0*-vaccinated (blue triangles) after challenge with EG.5.1. In contrast, sera from XBB.1.5-vaccinated and subsequently JN.1-challenged hamsters relocated to XBB.1.5 antigen. Altogether these results corroborate that XBB.1.5 vaccine-induced nAbs remain predominant in their specificity of the humoral response to newer Omicron subvariants. Importantly, in turn this also indicates that significant cross-reactive neutralizing responses against pre-Omicron variants (e.g. historical VOC Alpha) are not anymore induced by descendants of Omicron (e.g. XBB.1.5, EG.5.1, BA.2.86, or JN.1). Formally, the herein studied early and late SARS-CoV-2 variants may be considered representatives of two distinct serotypes.

## 4. Discussion

Alike for seasonal influenza, the WHO started issuing recommendations on how to regularly update current COVID-19 vaccines to keep up with the changing epidemiology and evolving antigenicity of emerging SARS-CoV-2 variants over time. Major vaccine suppliers are endorsed to adapt their products accordingly. In this study, we compared serological responses that are induced in hamsters by vaccination with Comirnaty® XBB.1.5 and subsequent experimental infection using the more recent, antigenically distant variants EG.5.1 and JN.1. Comirnaty® XBB.1.5 (Raxtozinameran) has been developed to match a particular Omicron subvariant dominating in Q1/2023 and has been rolled out since Q3/2023. Variant EG.5.1 has been on global rise since 7/2023, after which it has gradually been overridden by JN.1 gaining dominance since 12/2023 [2] (Figure 1a).

We show that vaccination with Comirnaty® XBB.1.5 induces in hamsters a strong neutralizing antibody response against its autologous antigen, XBB.1.5. However, despite cross-reactive binding antibodies being induced, nAb responses against the antigenically distant variants EG.5.1 and JN.1 were limited and waned rapidly over time. This appears to reflect similar patterns observed in humans where immune imprinting, shaped by earlier exposure to more ancestral strains of SARS-CoV-2, biases the response to newer variants [23]. These findings highlight how immunity to previous antigen encounter, while offering partial protection, may constrain the ability to generate de novo nAbs against significantly mutated variants like BA.2.86 or JN.1 encountered during secondary exposure, known as original antigenic sin [14].

Hamsters vaccinated with the monovalent XBB.1.5 mRNA vaccine showed strong protection against viral replication in the lungs when challenged with EG.5.1, corroborating that vaccine-induced immunity prevent lower respiratory tract disease. Lung damage was either minimal or entirely absent in hamsters exposed to the EG.5.1 and JN.1 variants. It remains unclear to what extent latter finding mirrors a reduced disease severity reported for humans infected with BA.2.86 and its descendant JN.1 [24]. Nevertheless, while these variants partially evade pre-existing antibody responses to cause significant morbidity in humans, they appear not to readily cause severe lung disease anymore in this experimental model. Importantly, our results are consistent with recent clinical observation showing that BA.2.86 and JN.1 infections mostly lead to mild illness in vaccinated individuals, despite their strong resistance to neutralizing antibodies [24]. Of note, our experimental model is limited to a single monovalent vaccination of naïve animals and single exposure, which is a sharp contrast to the human population exposed to several waves of SARS-CoV-2 variants, possibly multiple vaccine boosters and concurrently circulating multilineage SARS-CoV-2. Nonetheless, our model serves to illustrate the emergence and separation of SARS-CoV-2 serotypes due to gradual spike protein evolution (antigenic drift; e.g. B.1.1.7 – XBB.1.5 – JN.1).

Our findings underscore ongoing challenges in developing vaccines that can provide broad and durable protection against the continuously evolving cloud of co-circulating SARS-CoV-2 variants. The latest FDA-authorized COVID-19 vaccines are designed to target the spike protein of the KP.2 variant, one of the multiple descendants of JN.1 bearing three additional mutations (R346T, F456L and V1104L) [25]. Unlike JN.1, KP.2 was never dominant and was quickly surpassed by other variants, including KP.3.1.1 which by September 2024 represented the dominant variant in the US and Europe [20, 21]. In addition to the N-terminal domain deletion at S31, KP.3.1.1 differs from KP.2 in two other key spike protein positions at the receptor binding domain (i.e. Q493E, and reversion to R346 present in JN.1) [26], described to greatly decrease neutralizing activities of nAbs after JN.1 breakthrough infections [27] as well as therapeutic antibodies [28, 29]. The extent of antibody-mediated protection offered by a new KP.2-based vaccine is yet to be evaluated. Similar to influenza, prevention or even elimination of SARS-CoV-2 infection by vaccination remains a difficult-to-achieve task due to the high adaptability of the virus. Nonetheless, prior vaccinations with antigenically distant bivalent BA.4/5 or monovalent XBB.1.5 mRNA vaccine proved still efficacious in decreasing symptomatic and severe disease and hospitalizations, during periods which were already dominated by viruses of either newer Omicron XBB- or BA.2.86-derived sublineages [30].

Importantly, our data suggest that while nAb responses are essential for blocking infection, other immune mechanisms, such as T-cell-mediated immunity or Fc-mediated mechanisms, may contribute to the reduction of viral loads, as observed in the challenge with JN.1. Therefore, future vaccine designs should aim to induce broader and polyfunctional immune responses, encompassing not only humoral immunity based on nAbs but also involving other arms of adaptive immunity, to achieve durable protection against evolving SARS-CoV-2 variants.

In conclusion, our study provides further evidence that antigenic imprinting plays a dominant role in shaping humoral immunity against SARS-CoV-2 variants. While vaccines targeting ancestral or early Omicron strains offer partial protection, continuous updates and the inclusion of broader antigenic targets will be necessary to counteract the ever-evolving antigenic landscape of SARS-CoV-2.

## Author Contributions

Conceptualization, Y.A.A. and K.D.; methodology, J.D., E.M., K.G., Y.K., Y.A.A.; software, Y.A.A.; formal analysis, J.D., Y.A.A.; investigation, J.D., E.M., Y.K., B.W., Y.A.A.; resources, H.J.T., R.L., P.M., J.N., K.D.; data curation, J.D., Y.A.A.; writing—original draft preparation, J.D., Y.A.A., K.D.; writing—review and editing, ALL AUTHORS; visualization, J.D., Y.A.A.; supervision, Y.A.A., K.D.; project administration, Y.A.A., K.D.; funding acquisition, J.N., K.D. All authors have read and agreed to the published version of the manuscript.

## Data Availability Statement

Not applicable

## Acknowledgments

The authors are grateful to Carolien De Keyzer and Lindsey Bervoets for their support in challenge experiments; Tina van Buyten for providing the A549-hACE2-TMPRSS2 cells; and the staff of the animal facility at the Rega Institute (KU Leuven) for their excellent support.

